# Searching for a UV-filter in the eyes of high flying birds

**DOI:** 10.1101/336644

**Authors:** Malgorzata Zawadzka, Beatrix Ràcz, Dario Ambrosini, Carl Henrik Gørbitz, Jens Preben Morth, Alistair L. Wilkins, Anja Østeby, Katja Benedikte Prestø Elgstøen, Elsa Lundanes, Frode Rise, Amund Ringvold, Steven Ray Wilson

**Affiliations:** Department of Chemistry, University of Oslo, Post Box 1033, Blindern, NO-0315 Oslo, Norway; Enzyme and Protein Chemistry, Section for Protein Chemistry and Enzyme Technology, Department of Biotechnology and Biomedicine, Technical University of Denmark, Søltofts Plads, 2800, Kgs. Lyngby, Denmark; Faculty of Science and Engineering, The University of Waitato, Gate 1 Knighton Road, Private Bag 3105, Hamilton 3240, New Zealand; Department of Medical Biochemistry - Oslo University Hospital Rikshospitalet - PO Box 4950 Nydalen - NO-0424 Oslo, Norway; Department of Ophthalmology, Oslo University Hospital, Ullevål Postbox 4956 Nydalen NO-0424 Oslo

**Keywords:** Bird, Eye, UV, Liquid Chromatography, Nuclear Magnetic Resolution, Mass Spectrometry

## Abstract

An ultraviolet (UV)-absorbing compound of unknown identity is present in the aqueous humor of geese and other birds flying at high altitudes. A goose aqueous humor extract, that was believed to contain the UV protective compound which was designated as “compound X”, was fractionated and examined using a variety of spectroscopic techniques including LC-MS and high field one- and two dimensional-NMR methods. A series of compounds were identified but none of them appeared to be the UV protective “compound X”. It may be that the level of the UV protective compound in goose aqueous humor is much less than the compounds identified in our investigation, or it may have been degraded by the isolation and chromatographic purification protocols used in our investigations.

## Introduction

The eye is often compared to a camera; it forms an image of the environment on its photoreceptive layer, the retina. The diaphragm of the camera is analogous to the iris of the eye, and the lens focuses the light onto the most central part of the retina, the macula lutea. The light pathway through the eye media consists of cornea, aqueous humor, lens, and vitreous body. One decisive premise to achieve an optimal image is that all components of the media allow visible radiation (VR: 760-400 nm wavelength) to pass through. Shorter wavelengths (UV-A: 400-320 nm, UV-B: 320-290 nm, and UV-C: 290-200 nm) must be blocked to prevent phototoxic tissue damage, as the effect on biological tissue increases with decreasing wavelength. The most potent ultraviolet radiation (UVR) is largely absorbed by the ozone layer, minimizing damage on biological structures in general. In fact, only some 5% of all VR/UVR energy striking our planet is in the UV-range, and of this 97% is UV-A. However, it has long been known that this limited dose of actinic UVR reaching the ground is nonetheless able to cause harmful effects on the eye (1), and changes may occur at three different levels, i.e. the cornea (snow blindness/photokeratitis), lens (cataract), and retina (macula degeneration).

However, the human eye has some protection, e.g. the high levels of high molar absorptivity ascorbic acid present in the aqueous humor and corneal epithelium (30-300 times the serum level, respectively) (2–5). This view may explain why the ascorbic acid level in the corneal epithelium is high in diurnal and low/zero in nocturnal mammals (6), that the ascorbic acid content is much higher in corneal epithelium compared to aqueous humor in diurnal mammals, and that ascorbic acid content is higher in corneal epithelium of mammals living in the mountains, such as reindeer, compared to those living close to sea level (7). Furthermore, this hypothesis has also been supported experimentally as artificially increased levels of ascorbic acid in the aqueous humor of guinea pigs had a protective effect against UV-induced DNA damage to the lens epithelium (8), high ascorbic acid concentration protects the basal layer of the corneal epithelium by absorbing incident UVR (9), and ascorbic acid (as well as tocopherol) in the “aqueous humor” increased the viability of lens epithelial cells after UVR (10). There is also some evidence that systemic ascorbic acid supplementation used prophylactically may reduce the prevalence of hazy vision after photorefractive keratotomy (commercially available excimer lasers for refractive surgery are based on energy output at 213-193 nm wavelength) (11). Hence, ascorbic acid seems to act as “sunglasses” for the diurnal mammal eye. It would be natural to believe that the presence of ascorbic acid would be significant in the aqueous humor of high-flying birds, as these are exposed to vastly more UV radiation at e.g. 10,000 m altitudes. The aqueous humor is an integral part of the optic pathway of the eye media regardless of whether mammal or bird, and so an observation of low ascorbic acid content in avian aqueous humor was surprising (12).

What protect the bird’s eyes when flying at high altitudes, e.g. above the Himalayas (13)? Several UV-absorbing components, e.g. uric acid, tryptophan and tyrosine, have been proposed as being present in bird aqueous humor. In addition to these, a single, unknown component dominates the UV absorbing profile in goose aqueous humor, as observed with liquid chromatography-UV spectroscopy (LC-UV) (14). A lesser level of this compound is present in lower flying/nocturnal species e.g. duck and owl, and barely visible or absent in non-flying species (14). We hypothesize that this “compound X” has a protective role for the eye of birds and describe here our work towards identifying the unknown compound.

## Results and Discussion

### Liquid chromatography-UV analysis of highly polar substances in goose aqueous humor

Capillary liquid chromatography-ultraviolet absorption (LC-UV) 2D plots (**Figure 1**) shows chromophoric molecules (green frame), which were far more prominent in goose aqueous humor (top) compared to e.g. duck (bottom), resembling that observed previously (14). The green frame suggests the presence of multiple compounds. Compound 2 was however seen to form from compound 1 when samples was exposed to air, a clue that compound 1 (“compound X”) was sensitive to oxidation. The capillary LC system was coupled with electrospray-mass spectrometry (ESI-MS), a natural combination for gaining structure information of limited samples. Surprisingly, the molecules of interest had at this stage signals below the system’s detection limits, even when using high performance capillary LC-Orbitrap MS instrumentation.

**Figure 1.**
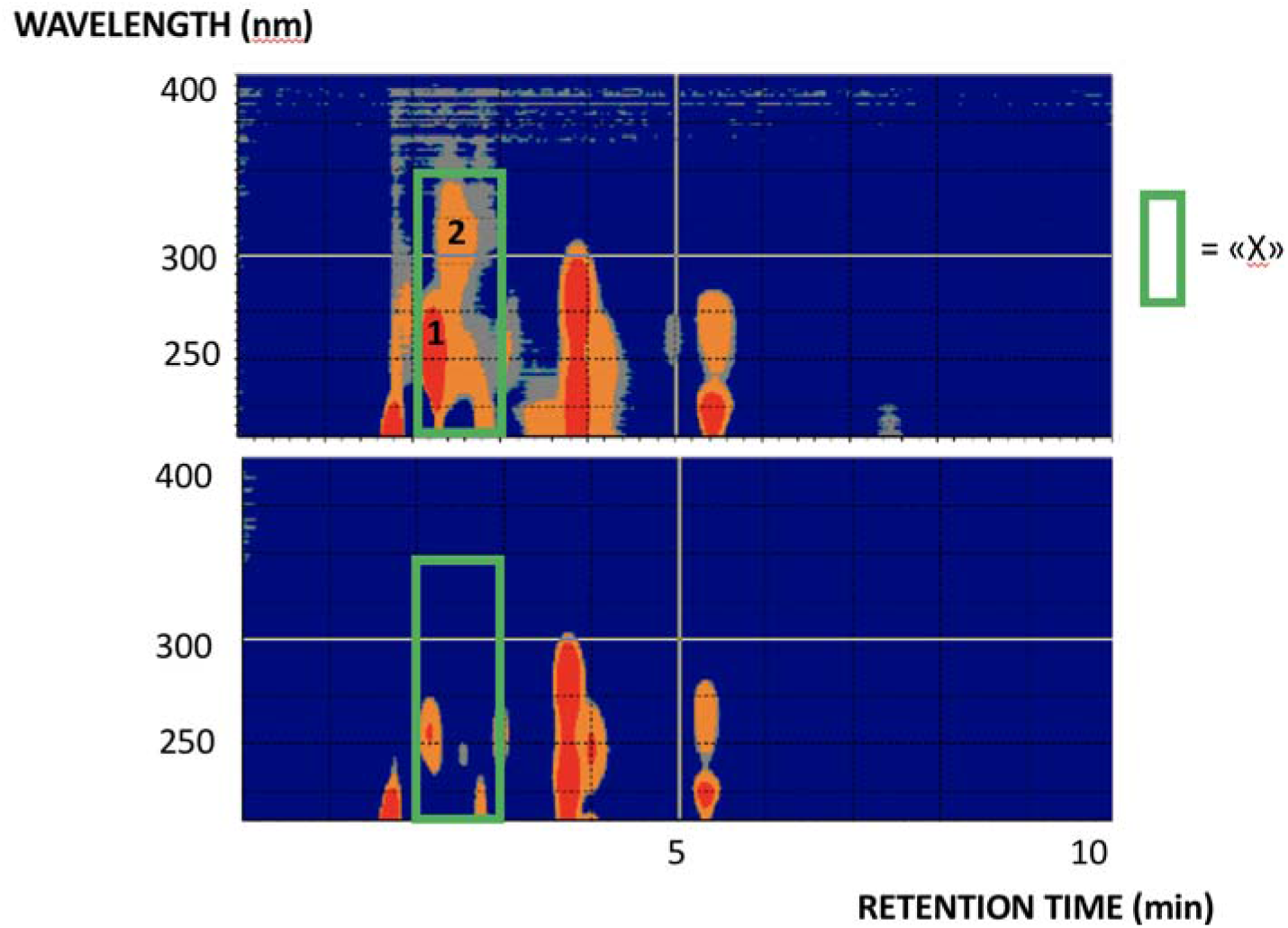
Two dimensional (capillary LC X UV) plots of aqueous humor from goose (top) and duck (bottom). Conditions are described in Materials section (Liquid chromatography and spectroscopy). 1 = “compound X”, and 2 is formed from “compound X”.

### Analysis of enriched material

To obtain improved signal intensities, “compound X” was enriched with semi-preparative scale LC for subsequent elucidation, using a modified RPLC column variant (ACE AQUA) for increased resolution of the polar analytes (**Figure 2,** 1D view; similar results were obtained using wild geese samples). Instant re-analysis of the isolated “compound X” revealed a pure peak, but the peak shape would be perturbed in subsequent analyses, again suggesting a reaction chemistry occurring when the analyte was chromatographed away from the aqueous humor. Moreover, when compound enrichment was attempted spending over 100 eye samples, a trace dark solid was produced. Size exclusion chromatography (SEC) could imply that the enriched material had (bio)macromolecular features (**Figure 3**). Hence, the analyte was assumed to form polymeric species/aggregates when concentrated and isolated from the aqueous humor. When treating “compound X” with trypsin, no enzymatic cleavage was observed, i.e. presence of tryptic peptides using MS analysis, which could rule out the presence of a protein (15). Crystallization procedures were subsequently attempted of isolated material, including developing an in-capillary crystallization procedure intended for limited samples (see **Supplementary Materials** for description of the procedure). However, only trace amounts of amorphous material were produced, unsuited for X-ray diffraction (XRD) experiments. ^1^H nuclear magnetic resonance (NMR) spectroscopy of the isolated material showed protons in the 6-8 ppm range (implying aromatic functionality), but again with insufficient sensitivity for structural analysis of the fractioned “compound X” peak.

**Figure 2.**
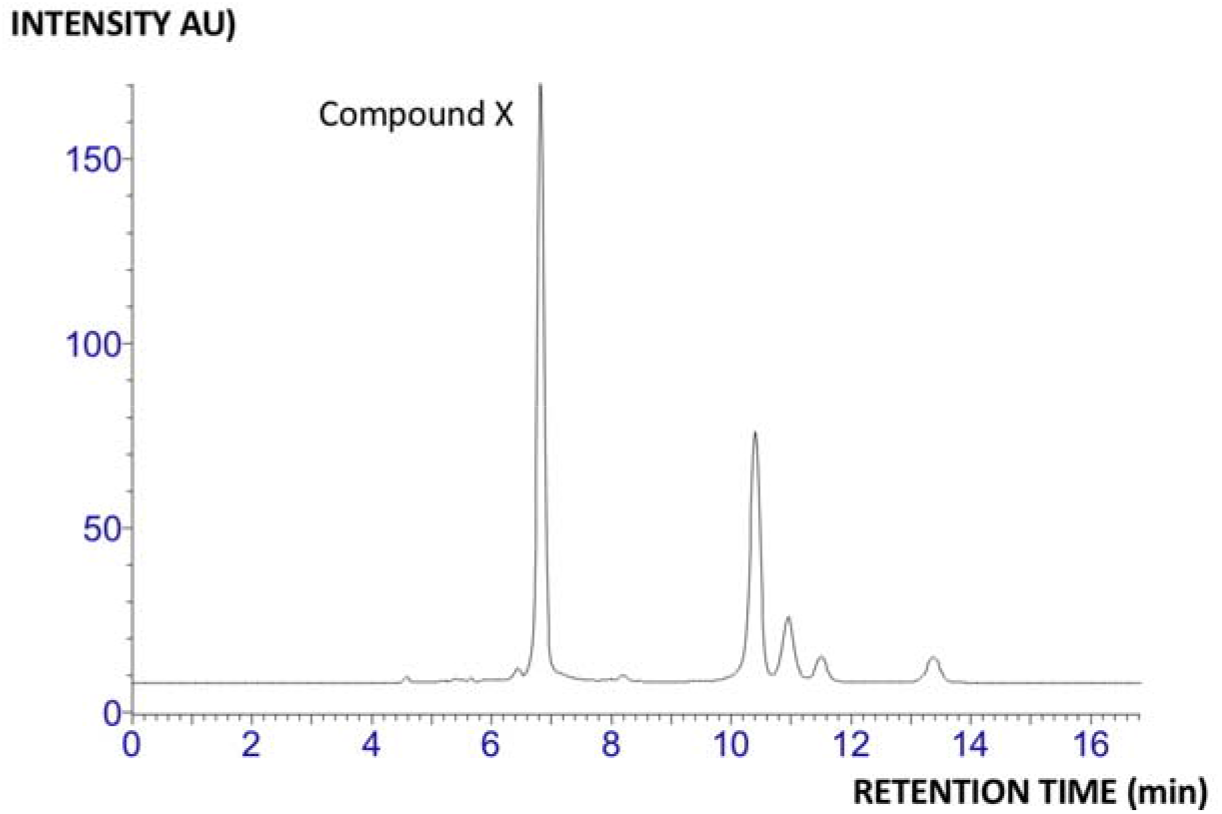
Reversed phase LC chromatography of goose aqueous humor, used for isolation of “compound X”. Conditions are described in Materials section (Liquid chromatography and spectroscopy).

**Figure 3.**
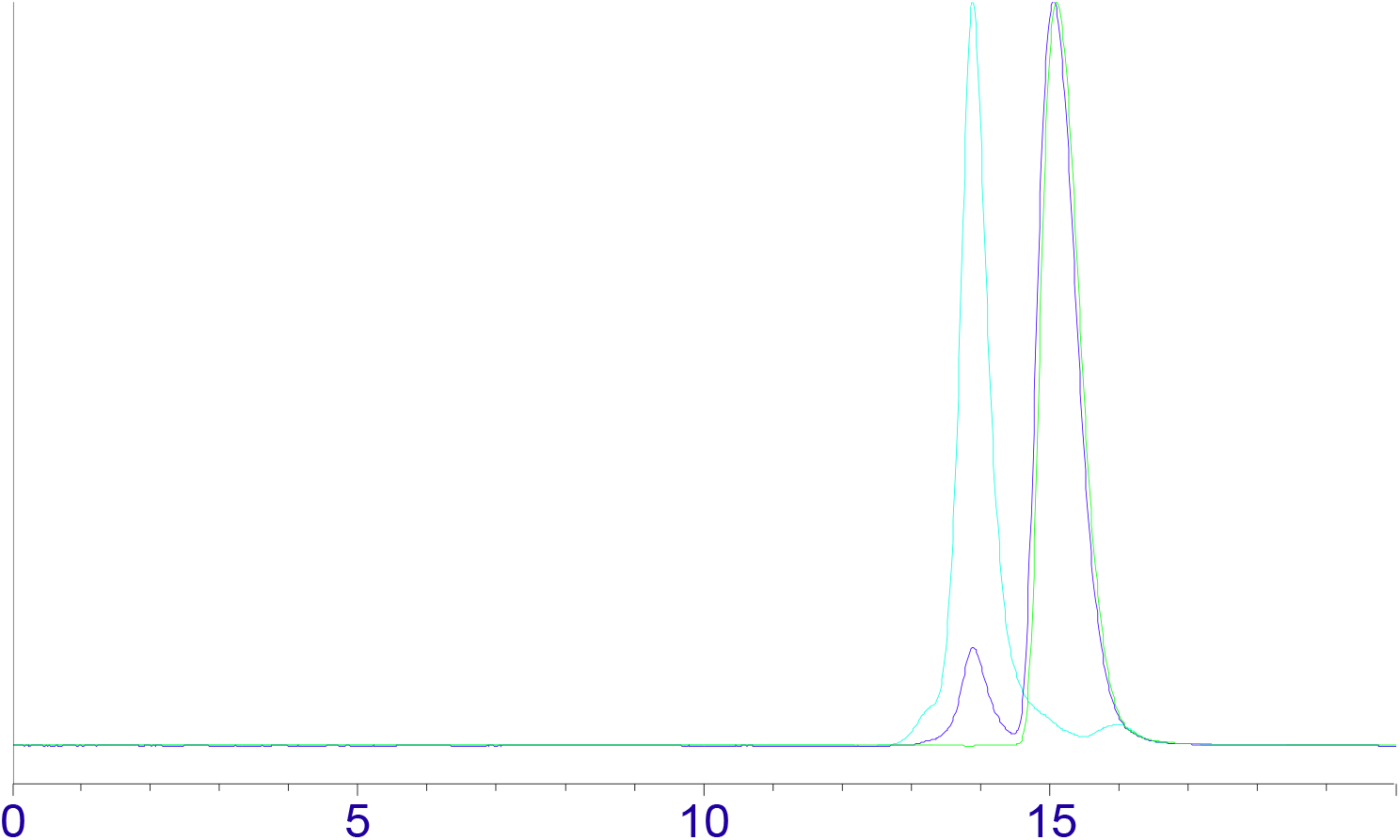
Size exclusion chromatography (SEC) of isolated “compound X” (light blue trace) and “compound X”/caffeine mixture (Vt marker, blue trace). Conditions are described in Materials section (Liquid chromatography and spectroscopy).

### Analysis of un-separated aqueous humor

The analytes were stable in capped aqueous humor, investigated with samples stored in a freezer, refrigerator, and subjected to freeze-thaw cycles, so analyses of un-chromatographed samples were undertaken. UV analysis confirmed that goose samples had elevated absorbance (**Figure 4, left**). Next, high field NMR (800 MHz) of non-chromatographed samples was performed, as this approach can be used for structural elucidation of un-separated molecules. The one dimensional ^1^H NMR spectra of un-chromatographed aqueous humor from various bird species goose, chicken, turkey resembled each other (Figure 4). Two-dimensional NMR spectroscopy enabled the identification several metabolites present, e.g. alanine, valine, leucine, isoleucine, tyrosine, phenylalanine, tryptophan and hypoxanthine. However, from inspection of the ^1^H spectrum and 2D nmr spectra we could not observe any aromatic substance with increased amounts in goose compared to other birds, in contrast to that observed with UV-based approaches (**Figure 4, right**). Unfortunately, the identity of any possible compound responsible for increased UV-absorption in goose samples was not established in our NMR experiments.

**Figure 4.**
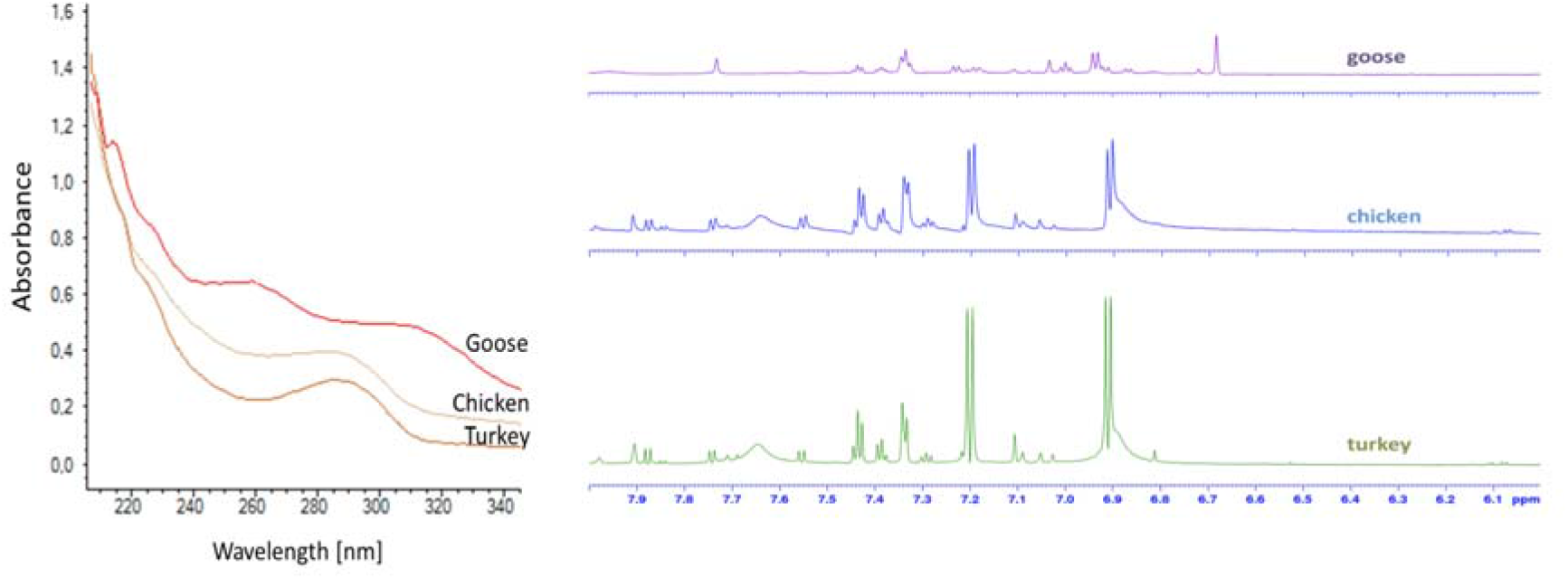
Left: UV absorbance of aqueous humor of goose, chicken and turkey. Right: 800 MHz NMR of the same samples. Conditions are described in Materials section (Liquid chromatography and spectroscopy and NMR Spectroscopy).

### Analysis of moderately enriched “compound X” fractions

The size exclusion chromatography experiments implied a polymerization occurring during intensive enrichments, so moderately enriched fractions of “compound X” were instead examined (5-10 fractions collected). Fluorescence spectroscopy experiments were performed, showing that the fraction had emitting properties (**Figure 5**). Revisiting mass spectrometry, now using conventional LC (2.1 mm ID column) coupled with an ESI source and a Q-Exactive mass spectrometer, the amino acid methionine was readily identified (also by the aid of an internal standard), see **Figure 6**. No other compounds in the fraction were detected/identifiable. However, we believe it is unlikely that methionine (MW 149.21 g/mol) is “compound X”, as this amino acid has few UV-absorbing properties. During our investigation we did speculate that the “compound X” fraction could have contained 5,6-dihydroxyindole (DHI), a monomer (MW 149.15 g/mol) of the biomacromolecule melanin (16, 17, 18), a hypothesis which could be supported by the indole-like properties and polymer-like features when performing isolation. Retention time comparisons with external standards were not possible, as commercially available DHI (Sigma) arrived in our lab as a partly polymerized/decomposed solid dark material (2 different batches). In addition, no indication of the presence of DHI was observed when NMR peaks were compared with those exhibited by another specimen of DHI which was not completely decomposed/polymerized. Furthermore, no NMR-signals in our isolated samples or in the pure eye liquid resembled nmr values reported in the literature (19). No HRMS peaks were clearly observed either on side of those arising from the M+ ion of methionine, effectively discounting the possibility that DHI was compound X.

**Figure 5.**
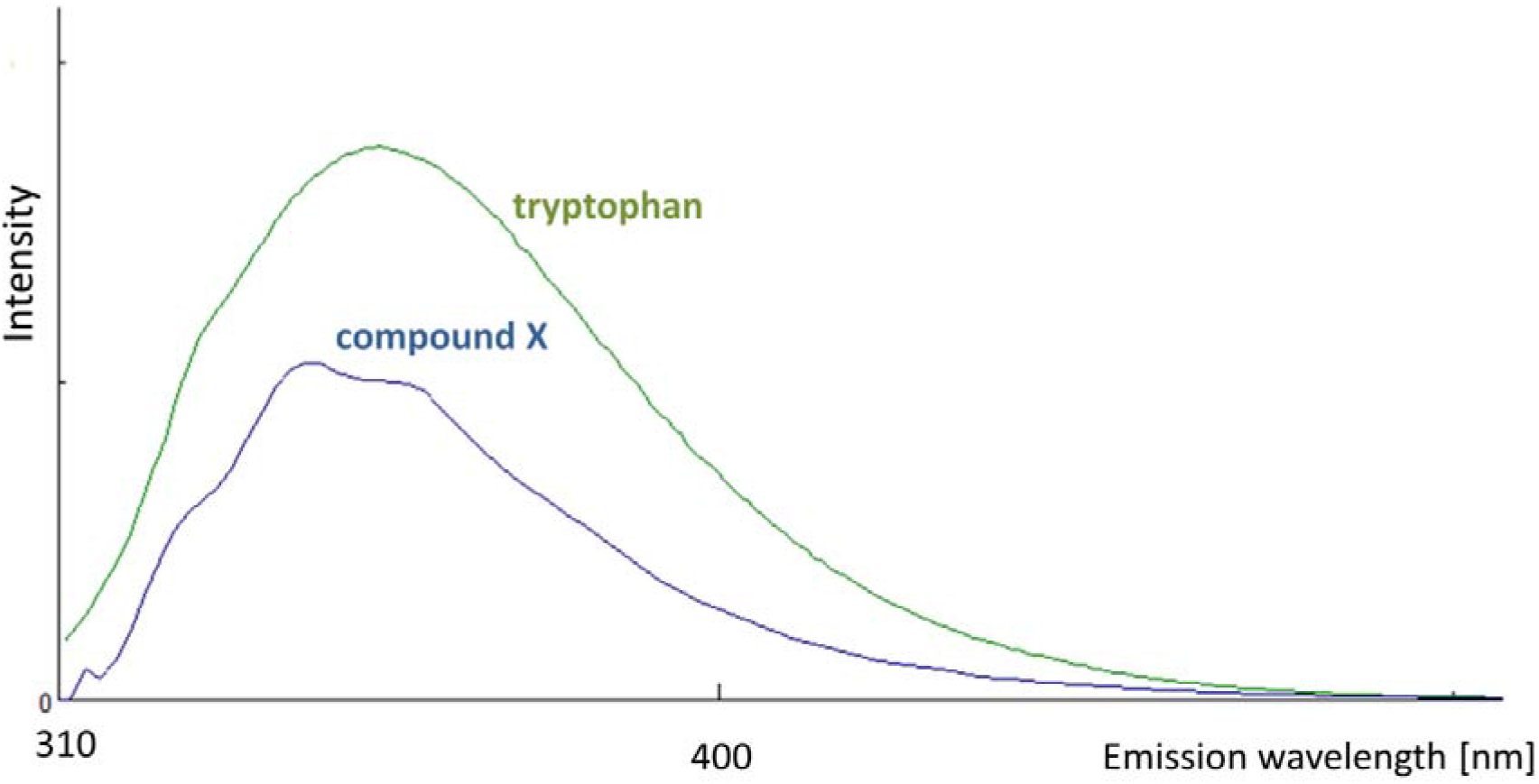
Fluorescence spectra overlay in 2D of “compound X” and tryptophan. Conditions are described in Materials section (Liquid chromatography and spectroscopy).

**Figure 6.**
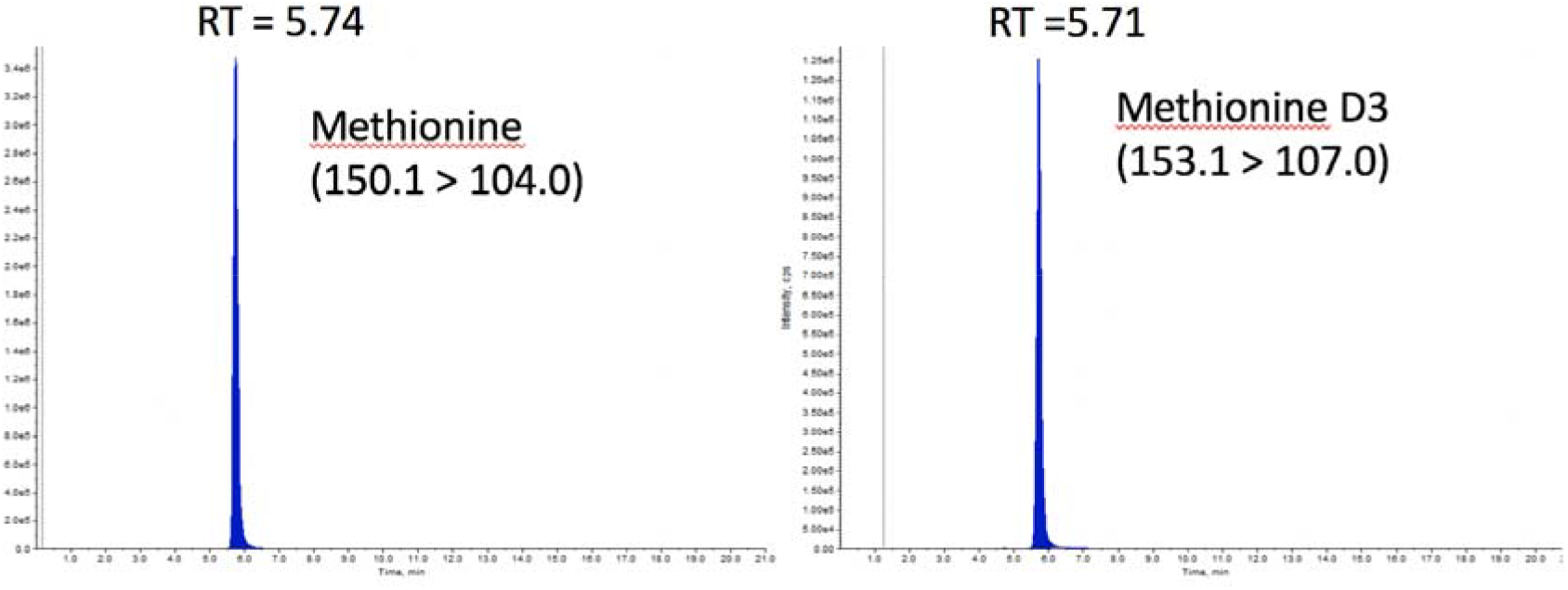
Methionine presence in fractionated “compound X” as observed with LC-MS and aided by an internal standard for retention time confirmation. Conditions are described in Materials section (Mass spectrometry).

## Conclusion

While a UV-absorbing compound is present in the eyes of birds the identity of the compound remains unknown. Although hundreds of samples were extracted, and analyzed by numerous techniques, the certain identification of this endogenous metabolite was not possible at this stage. This study serves as an example of the challenges that can be encountered when performing metabolomics of limited amounts analytes, and raises the possibility that degradation of the believed UV protective “compound X” may have occurred during the isolation and disappeared in our investigation.

## Methods

### Sample collection/preparation

Goose eyes (*Anserine*, White Italian) were made available to us each year in November, the only time when Holte Gaard (an ecological farm in Drangedal, Norway http://www.holtegard.no/) slaughter geese (for culinary purposes only). A veterinary would inspect the premises and routines the day of slaughter. Immediately after decapitation, aqueous humor was collected at the slaughterhouse through a limbal transcorneal canulation parallel to the iris diaphragm, 0.1-0.3 mL was obtained from each eye. Wild geese (Grey goose *Anser anser*, and Canadian goose *Branta Canadensis*) samples were donated from hunters. Chicken (*Gallus gallus*, Lomann), duck (Peking duck, *Anatinae*, English hybrid Cherry Waelly), and turkey (*Meleagris gallopavo*, British united turkey) were obtained under similar conditions as with the geese (farms: Prio Rakkestad and Revetal Vestfold, Norway).

### Liquid chromatography and spectroscopy

Capillary LC-UV was performed using a CapLC-UV system (Waters™, Milford, MA). LC analysis was performed with a PerkinElmer (Waltham, MA) series 200 LC pump chromatograph coupled to a Perkin Elmer series 200 autosampler and a Waters™ 486 Tunable Absorbance Detector set to a wavelength of 254 nm. The RP column was an ACE AQ HPLC column (ACE; 250 mm × 10 mm, 5 μm diameter particles, S/N A59810, Advanced Chromatography Technologies Ltd, Aberdeen, Scotland). The SEC column was a 4.6 × 300 mm TSKgel SuperSW3000 SEC column (Tosoh Corp., Tokyo, Japan).

The mobile phase used for RPLC was an isocratic mixture (99/1, v/v) of water and acetonitrile (with 0.1% addition of formic acid (FA) to both solvents for pH ≈ 2.7), with a flow rate of 5 mL/min. SEC separations were done isocratically with a flow rate of 0.35 mL/min and a mobile phase containing 0.05 M sodium phosphate and 0.3 M NaCl pH adjusted to 7.0. Injection loops of 100 μL and 1 mL were used. Thawed samples were injected without any pre-treatment. The smaller loop was pre-installed in the instrument’s autosampler; the bigger one was manually installed using a two-position six-port valve (Model 7000, Rheodyne LLC, Rohnert Park, CA, US). The use of 1 mL loop required manual injections. Chromatograms were obtained with the TotalChrom Navigator software version 6.2.1 (Perkin Elmer, Waltham, MA, US). Evaporation of manually collected fractions was conducted using a Speed-vac™ concentrator (SC110, Savant Instruments Inc., Hicksville, NY, US), in 1.5 mL vials (Thumbs up microtubes, Diversified Biotech, Boston, MA, US), with medium or no heating. Evaporating 1 mL of aqueous solution to complete dryness took about 6 h. Qualitative UV-vis analysis was performed on a Thermo Scientific NanoDrop 2000 UV-vis spectrometer. 2 μL sample were applied on the instrument’s pedestal. Type 1 water with 0.1 % FA was analyzed as a blank. The collected data were processed using the Chromeleon Chromatography Studio software (version 7.1.0.898, Dionex Corporation, Sunnyvale, CA, US). For fluorescence analysis an FP-8500 spectrofluorometer from Jasco with a Julabo F25-ED Refrigerated/Heating Circulator was employed. It was equipped with an ETC-815 Peltier thermostated single cell holder (water-cooled) for temperature control. The excitation wavelength was from 200 – 740 nm and emission wavelength from 305 – 750 nm without polarizing filters. Scan speed was 10000, temperature 25 °C. Data collection occurred with Spectra Manager (version 2.13.00). For measurements 350 μL of sample was pipetted into a quartz sample holder.

### NMR Spectroscopy

NMR spectra were aquired in pure aqueous humor liquid from geese with 10 % added D_2_O, 9: 1 (v/v) at 25 degree C using a Bruker AVIIIHD-800 spectrometer (Bruker BioSpin, Fallanden, Switzerland) equipped with a 5 mm TCI cryoprobe (^1^H, ^13^C, ^15^N) with automatic tuning and matching and Z-gradient accessories. ^1^H and 2D-HSQC spectra were recorded.

### Mass Spectrometry

With a Q-Exactive™ hybrid quadrupole-Orbitrap instrument from Thermo Scientific samples were transferred to mass analyzer using ESI in positive and negative mode with a spray voltage of 1.5 kV and a capillary temperature of 320 °C. Data were collected in a full scan mode (50 to 500 m/z and 150 to 2000 m/z). The pump flow was set to 0.02 mL/min (Fusion 100T pump, Chemyx Inc., Stafford, TX, US). For data collection and process control the Xcalibur software (version 3.0.63, Thermo Fisher Scientific Inc.) was used.

Quantification of methionine was performed using reversed phase ion pair liquid chromatography (LC) coupled to tandem mass spectrometry (MS/MS). The sample was prepared by adding isotopically labeled internal standard (IS) and was injected onto a reverse phase LC Uptisphere BP2 C18 column (50 × 2.1 mm, 3 μm, Interchrom©, Interchim, Montluçon, France) using a NexeraX2 LC system (Shimadzu, Japan). The mobile phase consisted of 1mM tridecafluoroheptanoic acid (A) and acetonitrile (B) and the gradient program were as follows: 0-1 min, 0-20% B; 1-6 min, 20% B; 6-9 min, 20-28% B; 9-16 min, 28% B. The flow rate was 200μL/min, the injection volume was 3 μL and the column temperature was 30⍰°C. The mass spectrometry analysis was performed on a Qtrap 4500 MD (AB SCIEX, Foster City, CA) using electrospray ionization in positive mode with the transitions 150.1 > 104.0 (methionine) and 153.1 > 107.0 (methionine-D3). All data collection and peak integration was performed using the Analyst^®^ software (version 1.6.2).

