## Supplemental Material for "Searching for a UV-filter in the eyes of high flying birds"

**Small-sample crystal growth by repeated heating and cooling cycles**

A single crystal X-ray diffraction (SCXRD) investigation of a new compound starts with crystallization, the goal normally being to obtain 0.1 – 0.8 mm sized crystals. In order to avoid precipitation of useless microcrystals as a result of massive nucleation from a supersaturated solution, it is imperative to gently transfer a compound´s solution from an unsaturated to a metastable state. Given time, a limited number of nuclei will eventually form, which can then grow by diffusion of additional material. The crystallization technique chosen will largely depend on the chemical properties of the compound of interest, mainly its solubility (Spingler *et al.*, 2012). A common feature for most substances is that solubility increases with temperature, implying that cooling of a hot saturated solution leads to precipitation and then also potentially crystals usable for SCXRD. Frequently, however, only microcrystals are obtained, implying that the process was either too rapid or that the compound itself has a challenging crystallization habit. In such cases the required specimens may be grown by applying repeated heating/cooling cycles (**Figure 1**). There has been a surprising lack of reports on instrumentation/hardware to enable this process, which appears mostly to be used for industrial processes (Kacker *et al.*, 2016; Jiang *et al.*, 2014), but also for proteins (Astier & Veesler, 2008; Zhang *et al*., 2008).

Our approach to crystallization with temperature programming is to use a setup consisting of a heating plate, an aluminum block that holds the test tubes/capillaries in an upright position, and an electric plug-in timer switch used for ON/OFF control of the plate (Figure 2a). An aluminum foil was wrapped around containers and capillaries to ensure more even heat distribution. The temperature on the plate was set to 70 °C and the ON/OFF cycles on the timer to 1 h/1 h. Temperature-changes on the plate were measured and read manually (Figure 2b**)**.

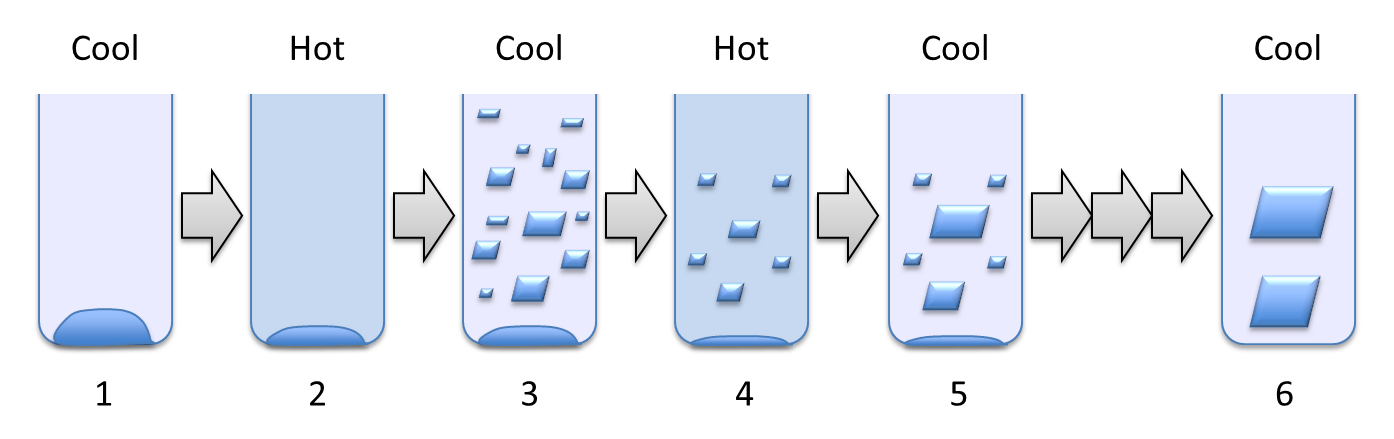

**Figure 1** Principles of crystallization by temperature cycling. The test tube or capillary is initially loaded with amorphous or microcrystalline material (1) that partly dissolves upon heating (2). Subsequent cooling leads to precipitation of microcrystals (3), which, with the exception of the largest ones, are dissolved in the next heating cycle (4). As the solution is now at equilibrium, few or no new nuclei are formed upon cooling, leaving the surviving crystals of the previous step to grow (5). Eventually more and more material is transferred to a small number of specimens, which after a number of cycles (triple arrow) reach a size suitable for SCXRD experiments (6).

The set-up was tested for a series of amino acids readily available in our lab. In terms of known crystallization habits, they range from very easy (Gly), through intermediate (Val and Ile) to more challenging (Leu). Each experiment typically being conducted for 84 cycles (7 days). Crystals grown in test tubes (Figure 2c) were harvested and mounted on the diffractometer in a regular manner.

A key focus of our work is to merge capillary liquid chromatography (cLC) systems (Rogeberg *et al.*, 2014) and SCXRD. cLC is highly suited for separating compounds in a mixture, especially if they are present in very limited amounts. Our intention is to trap separated compounds in capillaries in an “on-line” system, related to our previous coupling of LC and NMR (Mmatli *et al.*, 2007), and perform in-capillary SCXRD. Encouragingly, in-capillary crystal growth using heating/cooling cycles was possible (Figure 3a). Crystal-containing sections were attached to

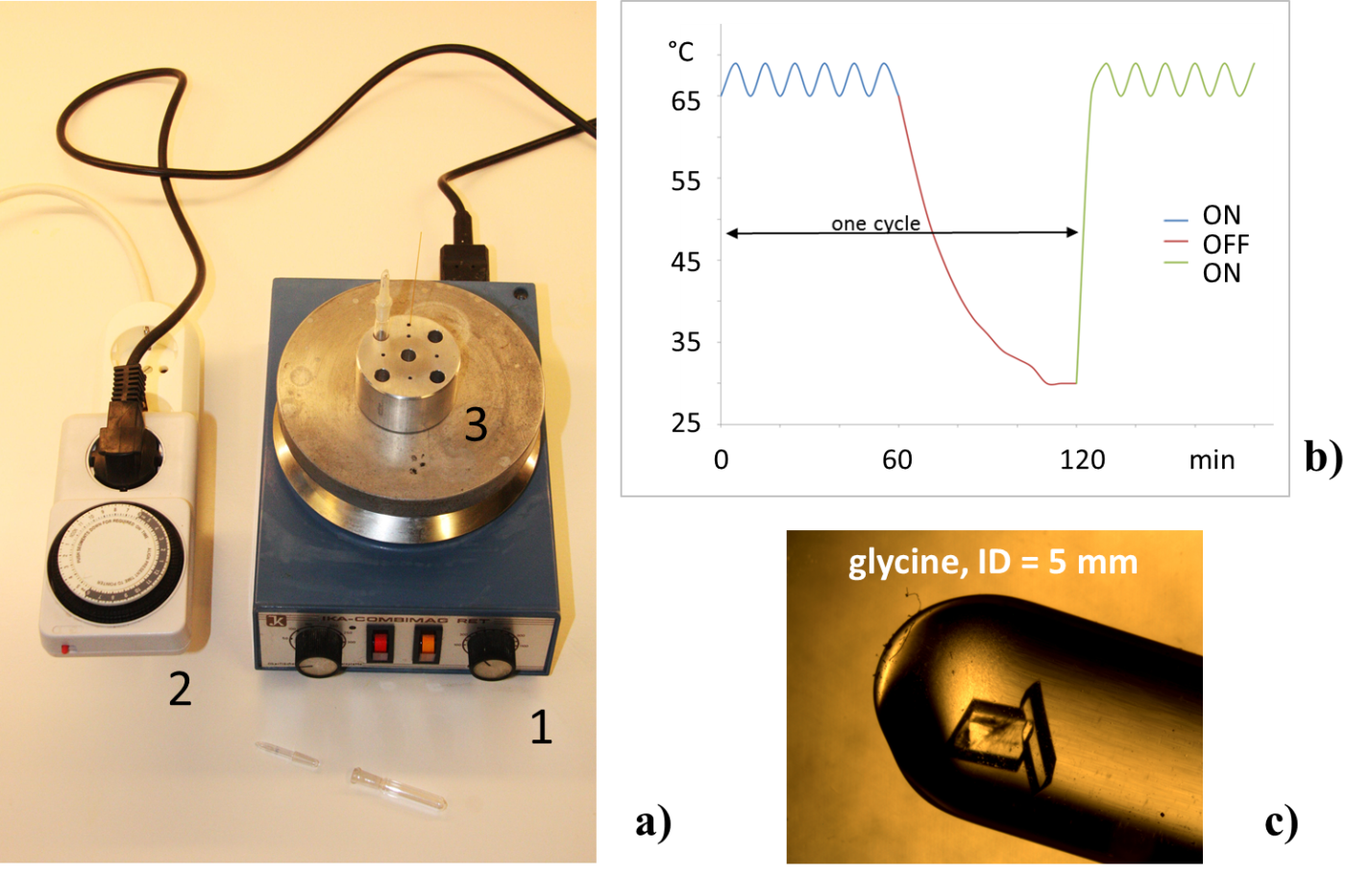

**Figure 2** (*a*) The set-up used consisted of a 620 W heating plate (1) connected a house-hold electric plug-in timer (max load 16 A, 3700 W), available at a cost of € 5-10 (2). Our aluminum block (3) is 5 cm in diameter with a height of 3 cm. It features five drilled holes for 7 mm diameter test tubes with stoppers (with ground joints, one set shown lying in front), and five for fused silica capillaries. (*b*) Illustration of the measured temperature during the temperature cycling. The temperature drops from about 68°C to 30°C when heating is off. Shorter cycles could have been used; a 30 min. cooling time would have reduced the temperature to about 35°C. (*c*) Example of crystals obtained using a temperature cycling program. The test tube was loaded with a few mg of microcrystalline glycine, filled with a methanol/water mixture (50/50, *v/v*) and capped. The top was finally wrapped in Parafilm to prevent the stopper from popping up during heating.

pin-heads, shock-frozen in liquid nitrogen to avoid formation of ice crystals, and mounted on the goniometer head. Satisfactory low temperature diffraction patterns were obtainable for capillaries with inner diameter 100 – 300 μm (Figure 3b), demonstrating a successful approach to SCXRD with minute amounts of sample (<< 0.1 mg).

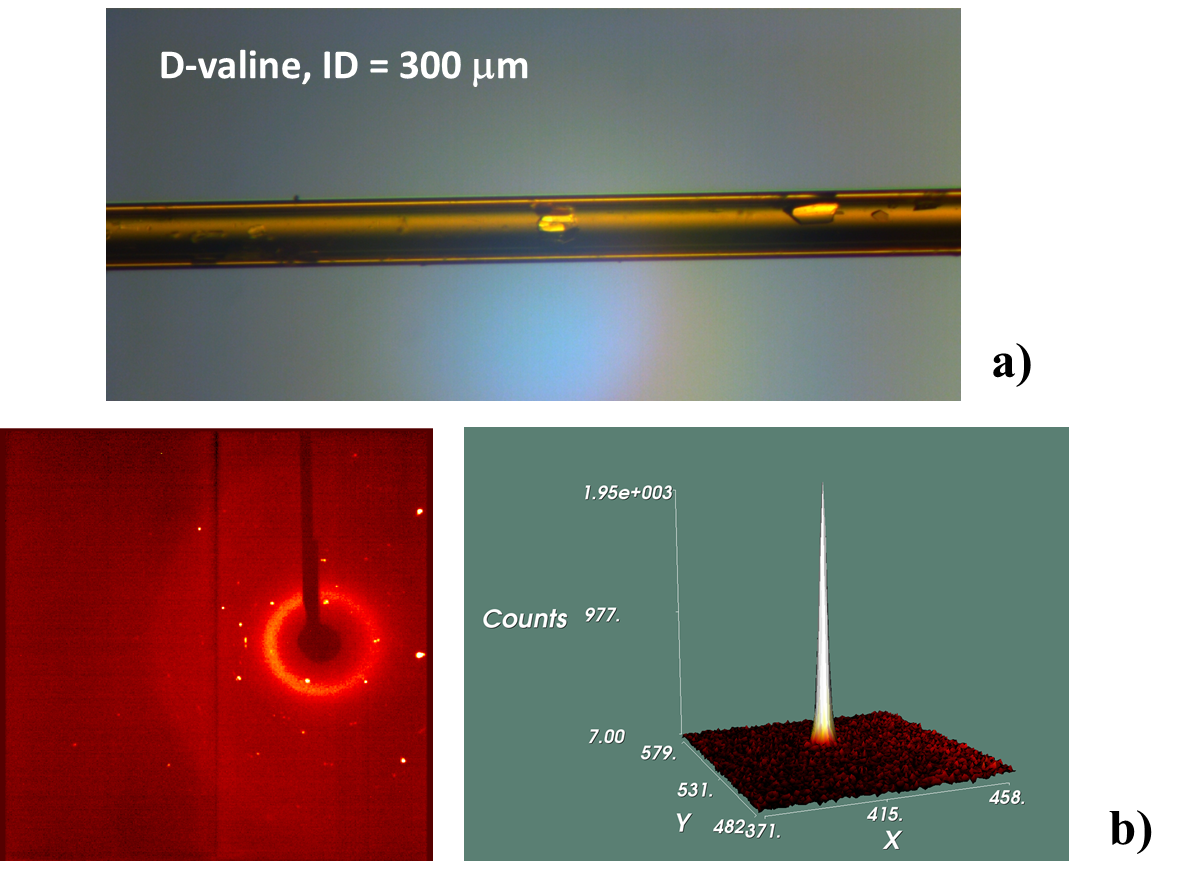

**Figure 3** (*a*) D-Valine crystals grown inside a 300 µm inner diameter (400 μm outer diameter) fused silica capillary. (*b*) Diffraction pattern from a 0.2 mm crystal inside a 4 mm capillary section attached to a pin-head. The quartz silica capillaries were sealed at both ends with epoxy glue drops (Araldite Rapid) or small silicone disks (Chrompack GC-septa). Diffuse scattering from the glass is noticeable, but did not affect the data quality. For capillaries with outer diameter 400 μm and inner diameter < 100 μm diffuse scattering became excessive.

We have seen that heating/cooling cycles for promoting single crystal growth for SCXRD is possible with simple means, allowing for e.g. in-capillary crystal growth with very small compound quantities.

SYNOPSIS

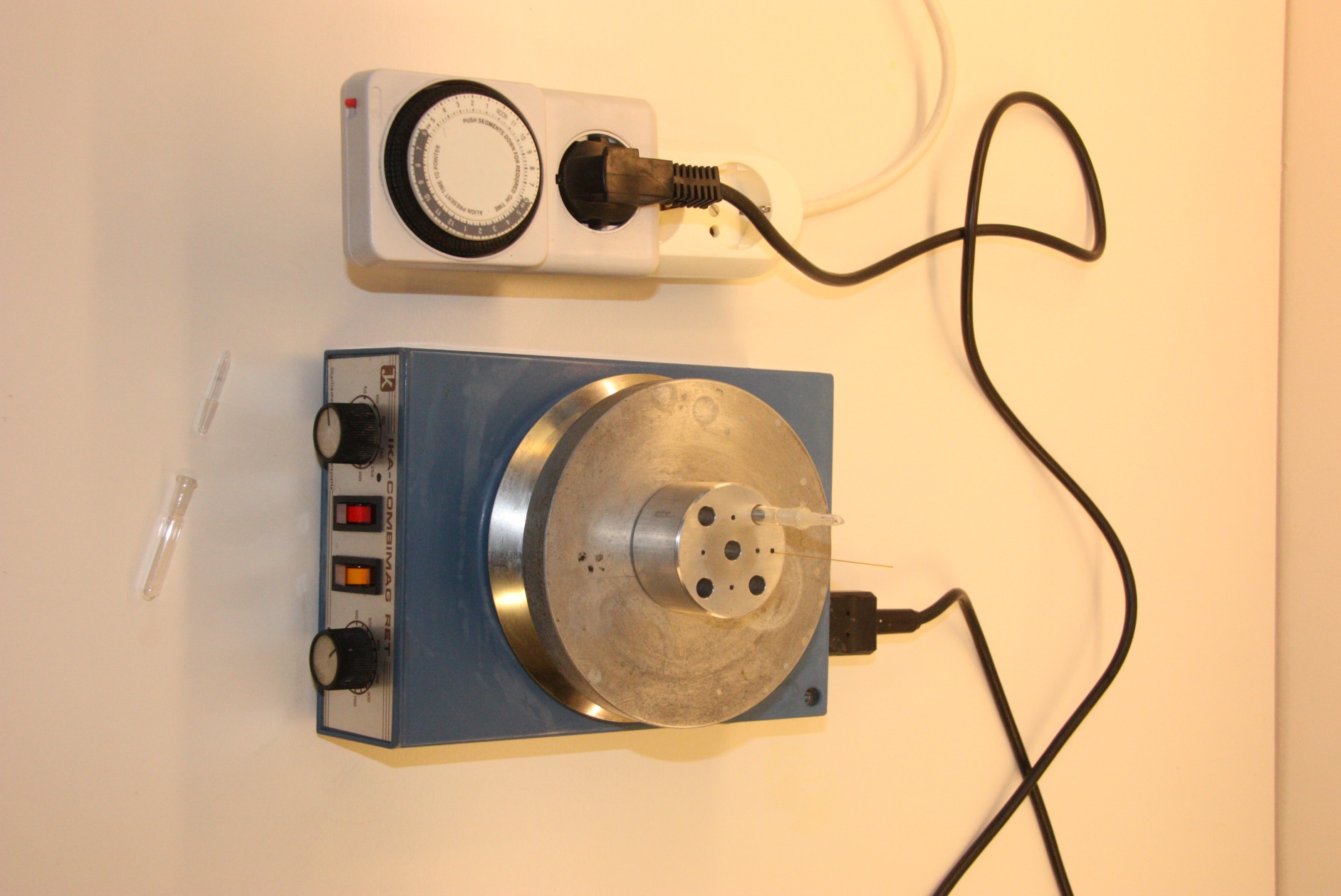

A very simple and cheap set-up for growing crystals of in test tubes or capillaries using a heating plate and a plug-in timer is presented. The method has the potential for extended use when crystals are difficult to obtain or in particular when the sample material is available in only sub-mg amount.
